# Transient pH changes drive vacuole formation in enzyme-polymer condensates

**DOI:** 10.1101/2025.01.16.633379

**Authors:** Nisha Modi, Raghavendra Nimiwal, Jane Liao, Yitian Li, Kyle J. M. Bishop, Allie C. Obermeyer

## Abstract

Intracellular membraneless organelles formed by the phase separation of biomolecules are essential for cellular functioning. These biomolecular condensates often exhibit complex morphologies in response to biological stimuli. *In vitro* condensate models help elucidate the mechanism of formation and the associated function of these hierarchical assemblies. Here, we use an *in vitro* model to investigate the formation of hollow internal regions, or vacuoles, within the condensate interior in response to a pH change. Our experimental system is a pH-responsive complex coacervate formed by the anionic glucose oxidase enzyme phase separating with the weak polycation, DEAE-dextran. Fast rates of pH decrease and larger droplet sizes trigger vacuole development within the coacervates. We show that the emergence of vacuoles is a non-equilibrium process caused by the diffusion-limited exchange of condensate components during a fast pH change. We develop a theoretical model that captures how a phase-separating system responds dynamically to changes in system conditions, particularly pH. Our qualitative phase diagram aligns with our experimental results, showing that rapid pH changes shift the phase boundaries, triggering spinodal decomposition and inducing vacuole formation within the condensates. Our pH-sensitive *in vitro* coacervate model provides a platform to modulate the internal structure of ternary phase separating systems and gain insights into the mechanisms controlling condensate organization *in vivo*.

## Introduction

Biomolecular condensates exhibit diverse structures and perform critical cellular functions. Biomolecules condense or phase separate in cells to create distinct microenvironments that enable a range of cellular functions from signaling [1] to cell division [2]. These condensed phases are not always uniform in composition, often displaying features like multiphase or layered structures, including vacuoles [2], [3]. These hierarchical structures have been shown to be important for certain biological functions. A classic example is the nucleolus, which is a three-layered condensate, wherein each immiscible layer is involved in a specific stage of ribosome biogenesis, from rRNA transcription to rRNA processing, and ribosome assembly [4]. Preferential partitioning of miRNA within the inner layer of multilayered cytoplasmic Processing Bodies (PBs) is coupled to their role in translational regulation [5]. Heterogeneous condensate structures can also facilitate their formation and dissolution in response to stress [6]. Understanding how, when, and why these higher order structures form has largely yet to be determined.

*In vitro* or synthetic models of condensates can be useful for gaining mechanistic insight into the assembly and function of biomolecular condensates, including those with heterogeneous structures. The immiscibility of each of the nucleolus layers is now attributed to differences in surface tension elucidated by *in vitro* experiments on condensates composed of endogenous nucleolar proteins [4]. Fluorescence recovery after photobleaching (FRAP) experiments performed on synthetic condensates composed of PGL-3 and MEG-3 proteins found in P granules show that the heterogeneous condensate structure is key to maintaining the concentration gradient of P granules during the development of *C. elegans* embryos [7]. Fully synthetic *in vitro* models of condensates have also been shown to undergo structural changes, including the formation of vacuoles within the condensate interior, wherein the composition of vacuoles resembles the external dilute phase. This internal reorganization can be driven by active processes [8], [9] or by altering external stimuli. For example, the appearance of hollow vacuoles within synthetic polyA-PEG condensates is regulated by temperature changes, and results from the slow mobility of condensate components during rapid temperature shifts [10]. Similarly, exposing initially homogeneous soy glycinin condensates to changes in temperature, or salt or pH, leads to a transition to hollow condensates [11]. These changes in external stimuli modify the hydrophobic and electrostatic interactions dominating the soy glycinin condensate such that the structural ordering of condensate components at the surface of the hollow condensate is more favored.

As exemplified by the investigation of soy glycinin, changes in electrostatic interactions are crucial to condensate structure, but they are relatively under explored in the context of condensate structuring. Electrostatic interactions between biomacromolecules play a key role in the formation of these condensates, and several biological processes, such as post-translational modifications and subcellular proton gradients, fine-tune these interactions [12], [13]. Analogous to cellular control over condensate assembly *in vivo*, properties of *in vitro* condensate models can be modulated by adjusting the electrostatic interactions between the phase-separating components. In this paper, we focus on vacuole formation in protein-polymer condensates regulated by pH, a biologically relevant variable. Notably, the presence of pH gradients of magnitude greater than 0.5 pH units have been observed within biomolecular condensates [12], [13], highlighting the important role that pH plays in condensate formation, structure, and potentially in biological function.

Herein, we demonstrate that vacuole formation in pH-sensitive coacervates is a non-equilibrium process brought about by the diffusion-limited ability of the coacervate components to equilibrate with the external pH change. Our pH sensitive *in vitro* model consists of complex coacervates composed of the enzyme glucose oxidase and the polycation DEAE-dextran. We develop an experimental method to systematically vary and quantitatively track pH changes around droplets using optical microscopy. Our experimental results, supplemented with a qualitative model, show that the slow mobility of the phase-separating components across the droplet boundary during a fast pH drop results in the coacervate entering the spinodal region. We discuss how competition between the rate of change in external stimuli and time scales required for a phase-separating system to relax to equilibrium can provide a mechanistic understanding of vacuole formation in such associative coacervate-type condensates.

## Results

### Complex coacervates form vacuoles upon rapid pH change

The equilibrium phase behavior of oppositely charged polyions, or complex coacervation, is known to depend on a range of parameters that influence electrostatic interactions between the polyions. This primarily includes the balance of opposite charges on the polyions, which is a function of both the composition and pH for weak polyelectrolytes. While it is intuitive that complex coacervates should be sensitive to the solution pH, it is not clear what happens to a coacervate when pushed out-of-equilibrium by a change in the pH. Our model system of glucose oxidase and DEAE-dextran has previously been shown to respond to changes in solution pH [14]. Here we probe the equilibrium and out-of-equilibrium phase behavior of this system to answer the question: what happens to a coacervate when the pH is changed?

To investigate how complex coacervates respond to changes in pH, we first established the equilibrium phase behavior as a function of pH. We focused efforts on basic pHs (8-10), as this range includes the pKa of the polycation (pKa ~ 8.8) [14]. We then examined the dynamics of these coacervates when driven out of equilibrium by the addition of acid, which creates a charge imbalance within the system.

Initial turbidity experiments show a dependence of coacervate formation on pH and mixing ratio of the participating polyelectrolytes. At pH 10, mixing the anionic enzyme (estimated charge ~ −90) with the cationic polymer (estimated charge ~ +400) results in phase separation with the turbidity maximum occurring at approximately a 3:1 ratio of enzyme to polymer (0.76 mass fraction of the enzyme). As pH decreases from 10 to 8, the negative charge on the enzyme decreases from ~ −90 to ~ −60, and the positive charge on the DEAE-dextran increases from ~ +400 to ~ +700. The change in the charge on these weak polyelectrolytes as a function of pH alters the equilibrium phase behaviour and ultimately the composition of the enzyme/polymer coacervates. As the pH decreases, the turbidity maximum shifts to a higher fraction of glucose oxidase to balance the increase in positive charge on the polycation; the turbidity peak shifts to a 5.25:1 ratio of enzyme to polymer (from 76% glucose oxidase to 84% glucose oxidase). At a decreased pH ~ 8, coacervate formation is initially not observed with the lower percentage of glucose oxidase; however, micron-sized droplets gradually emerge over extended time scales. Optical microscopy shows the presence of coacervate droplets with uniform interiors at both pH 10 and ~ 8 at a 0.76 mass fraction of the enzyme.

Driving coacervates at pH 10 out of equilibrium by adding acid can lead to the formation of vacuoles (Fig. 1). A decrease in pH induces a charge imbalance, altering the equilibrium composition of the dense and dilute phases. Under some conditions, the addition of acid to coacervate droplets initially at pH 10 results in the emergence of hollow regions within the droplets. We hypothesized that when the pH decreases from 10 to 8, smaller coacervate droplets or those exposed to slow rates of pH decrease would remain homogeneous and resemble equilibrium droplets at pH ~ 8 (Fig. 1a). In contrast, vacuoles would form within larger coacervates or droplets subjected to rapid pH decreases, with the composition of vacuoles resembling that of the dilute phase (Fig. 1 a and b). To program vacuole formation in coacervates, it is essential to understand how droplet properties and external perturbations influence the onset and dynamics of vacuole formation. To achieve this, we designed an experimental setup that allows us to vary the rate of pH change while simultaneously studying how droplet properties evolve as a function of the surrounding pH.

**Figure 1:**
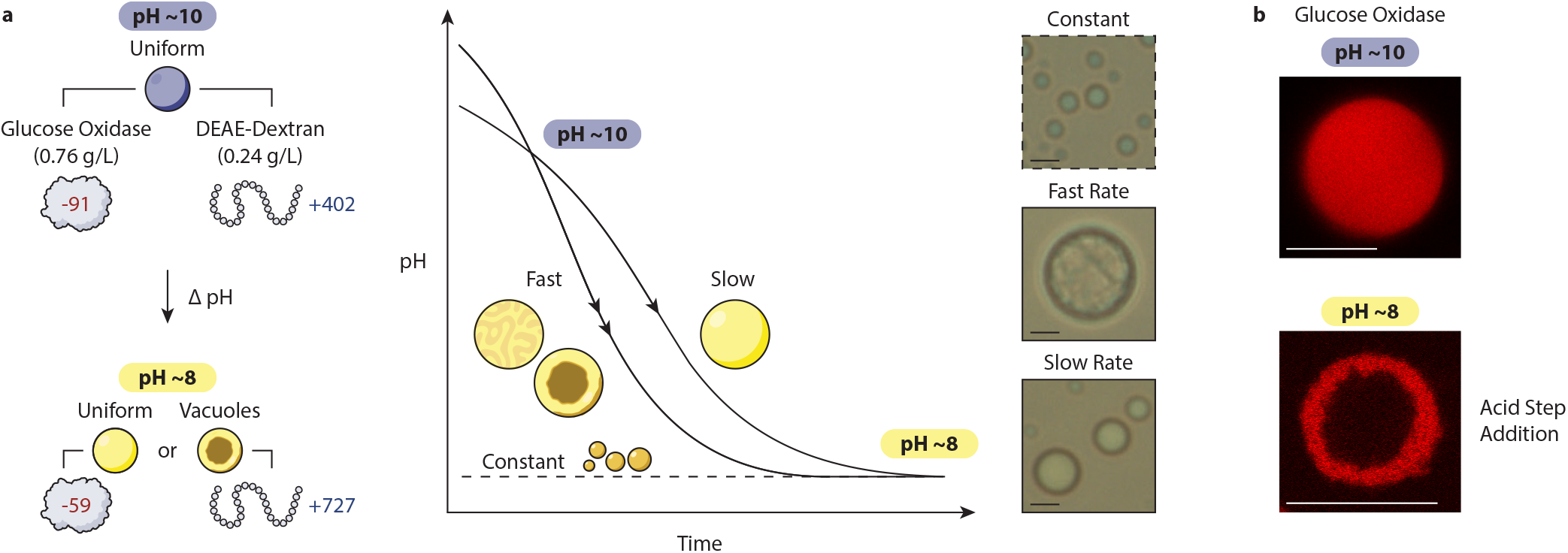
Complex coacervates of glucose oxidase and DEAE-dextran show path-dependent vacuole formation upon a decrease in pH. (a) The anionic enzyme glucose oxidase phase separates with cationic DEAE-dextran to form uniform condensates at pH 10. Depending on the rate of pH decrease, uniform condensates initially at pH 10 may or may not form vacuoles at pH 8. (b) Confocal microscopy images of glucose oxidase labeled with Alexa Fluor 594 and unlabeled DEAE-dextran. Vacuole formation occurs upon the addition of acid as the pH is lowered from 10 to 8. Scale bars are 10 µm.

### Capillary experiments generate spatiotemporal pH profiles

To investigate the conditions under which vacuoles form, we first designed an experimental setup that subjects coacervate droplets to varying spatiotemporal pH profiles in a controllable and reproducible manner. The setup allows us to observe morphological changes in the droplets and quantify pH changes *in situ*.

Our experimental setup consists of a coacervate solution contained in a long, thin glass capillary with a source of acid at one end (Fig. 2a). The acid source is a small agarose gel loaded with acid. As acid diffuses from the gel into the coacervate solution, the pH of the coacervate solution decreases. This configuration allows us to model the diffusion of acid across the capillary length as a onedimensional reaction front and obtain varying rates of pH decrease at different distances from the gel interface [15]. To prevent the wetting of coacervate droplets on the capillary walls, the interior surface is passivated with mPEG-silane. The coacervate solution is prepared by mixing glucose oxidase and DEAE-dextran solutions in water adjusted to pH 10 at a mixing ratio corresponding to the peak turbidity (0.76 mass fraction of enzyme). As the acid front travels across the length of the capillary, coacervate drops closer to the gel boundary experience a more significant and faster drop in pH, while drops further away from the gel are exposed to a slower and smaller drop in pH. Using this setup, coacervates of varying sizes are exposed to different rates and magnitudes of pH decrease depending on the distance from the gel boundary.

**Figure 2:**
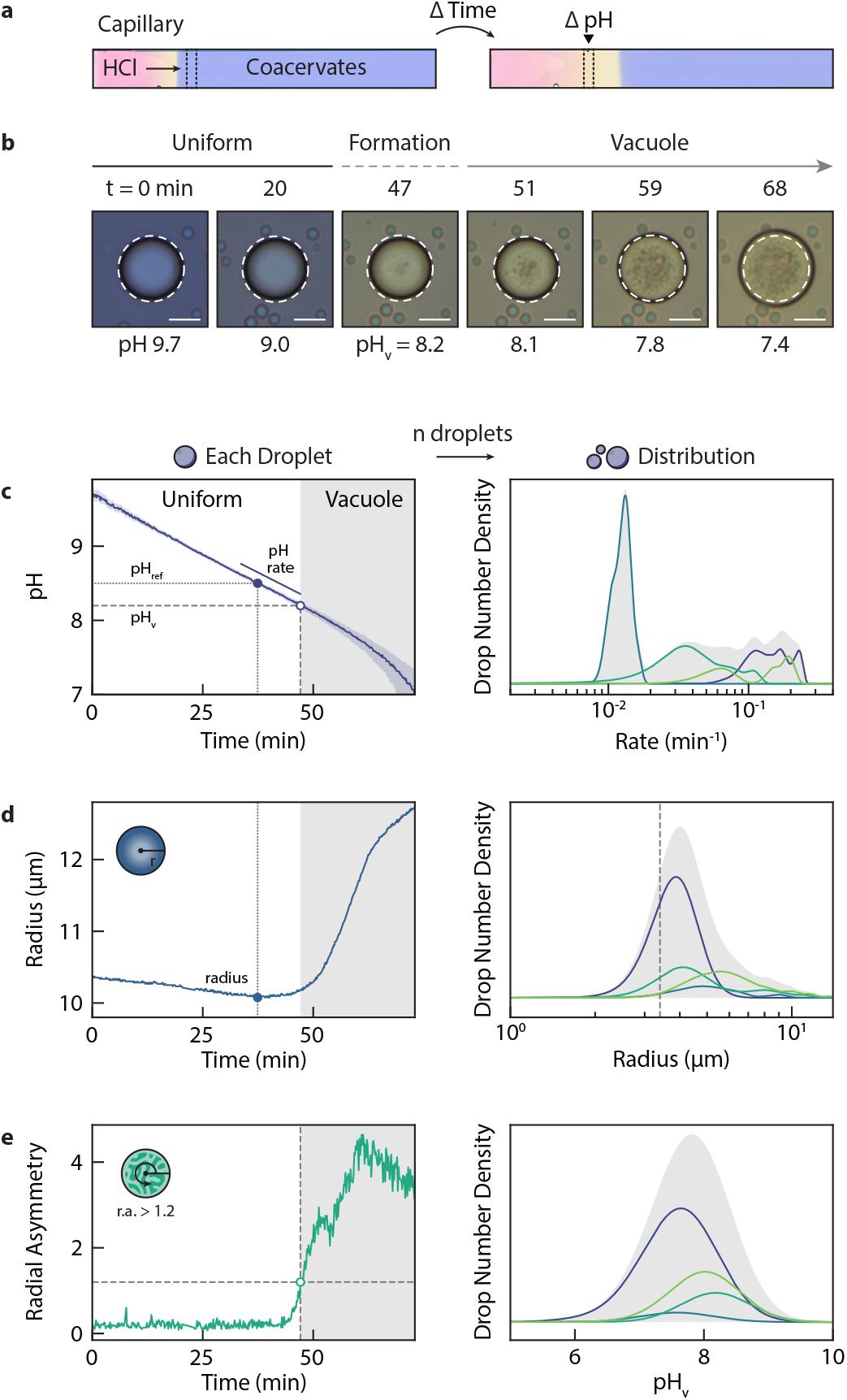
Capillary chamber can be used to generate a distribution of pH profiles. (a) Representative images of a capillary experiment. The capillary contains an agarose gel with HCl on one end, while the rest of the capillary is filled with a coacervate solution in water at pH 10. Both acid and coacervate solutions contain 0.2 mg/mL of Xylenol Blue, a pH indicator enabling spatiotemporal visualization of the pH. The total macromolecular concentration of the enzyme and DEAE-dextran is 1 mg/mL, and the mass fraction of glucose oxidase is 0.76. (b) Microscopy images of vacuole formation in a representative droplet over time as the pH decreases. Scale bar is 10 */mu*m (c) (left) The instantaneous pH of a single droplet can be monitored over time by analyzing the pixel color (Supplemental Note 2). The rate of pH change experienced by a single droplet is estimated by the slope at pH 8.5 (solid line). (right) Imaging at different distances from the gel front enables a distribution of rates of pH change to be visualized (d) (left) The instantaneous radius of a single droplet can be quantified over time. For comparison, drop radius is defined as the radius at pH 8.5. (right) Droplets of varying size were imaged. Only drops with a radius greater than 3.4 µm were further analyzed for vacuole formation. (e) (left) The radial asymmetry of a drop was also quantified from images and used to automatedly detect vacuole formation. The time at which the radial asymmetry was greater than 1.2 was used to determine the pH of vacuole formation (*pH*_*v*_). (right) *pH*_*v*_ varied for the *>*200 drops analyzed.

We use the colorimetric pH indicator Xylenol Blue to effectively quantify the pH of the solution in the capillary. With a pKa of 8.94, Xylenol Blue is sensitive to pH changes in the desired range (10–8). As the pH decreases from 10 to 8, Xylenol Blue undergoes a color change from blue to yellow. The color of the indicator, captured in microscopy images as RGB values, can be converted to pH using the relationship between the fraction of deprotonated indicator (*δ*) and the color coordinates in the CIE XYZ color space (Supplementary Note 2). While the pH in the capillary may drop below 7, the accuracy of the pH measurements diminishes as the pH deviates further from the pKa of the indicator. Therefore, we limit our analysis to the range of pH 10–7, where the indicator provides reliable measurements.

We evaluate the change in morphology of individual coacervate drops as a function of their temporal pH profile using microscopy and image analysis. Each droplet is tracked and analyzed to extract parameters such as the pH, drop radius, and internal inhomogeneities in the form of vacuoles. The Radial Variance Transform (RVT) package in Python [16] is used to identify droplet centers, and radial intensity variations are analyzed to detect vacuoles within otherwise homogeneous droplets. The rate of pH decrease is calculated by fitting a linear model to the pH profile, focusing on the region centered at pH 8.5, as most vacuoles form below this pH (Fig. 2c).

By observing an individual droplet within the capillary, we can track changes in the surrounding pH, radius, and internal structure over time, providing detailed insights into the droplet’s response to spatiotemporal pH changes (Fig. 2b). For an example droplet, the droplet begins as a homogeneous sphere at pH 10. As the pH decreases, indicated by a color transition from blue to yellow, the droplet shrinks in size (Fig. 2d). Under specific conditions of pH decrease, vacuoles form within the droplet, accompanied by swelling. The onset of vacuole formation is characterized by an increase in radial asymmetry (Fig. 2e). A radial asymmetry value exceeding a threshold of 1.2 is used to define vacuole formation, with the corresponding pH (pH_v_) and radius at this point marking the critical conditions for the transition.

Analysis of data from four independent capillary experiments includes rates of pH decrease spanning an order of magnitude (0.01 to 0.24 min^−1^, Fig. 2c) and measurements of droplet sizes ranging from 3.4 to 13.7 microns (Fig. 2d). We selectively analyze droplets with radii greater than 3.4 microns because the number of pixels in droplets smaller than this minimum size is inadequate for accurate image analysis. The results show that vacuole formation within coacervates occurs at pH values below 8.5 (Fig. 2e).

### Onset of vacuole formation depends on drop size and rate of pH change

Our results show that the onset of vacuole formation depends on the rate of pH decrease and the drop size (Fig. 3c). Using the capillary setup, imaging within a single field of view allows simultaneous monitoring of tens of drops of varying sizes, all experiencing similar rates of pH change (Fig. 3d). Among coacervate drops with identical spatiotemporal pH profiles, larger drops form vacuoles at slower rates of pH decrease and at higher pH values. For example, Fig. 3a compares the change in morphology in two drops of different sizes subjected to the same rate of pH decrease, from pH 9.4 to 8.3, over approximately 15 minutes. While the larger drop exhibits vacuole formation at pH 8.5, the smaller droplet does not form vacuoles until pH 8.3.

**Figure 3:**
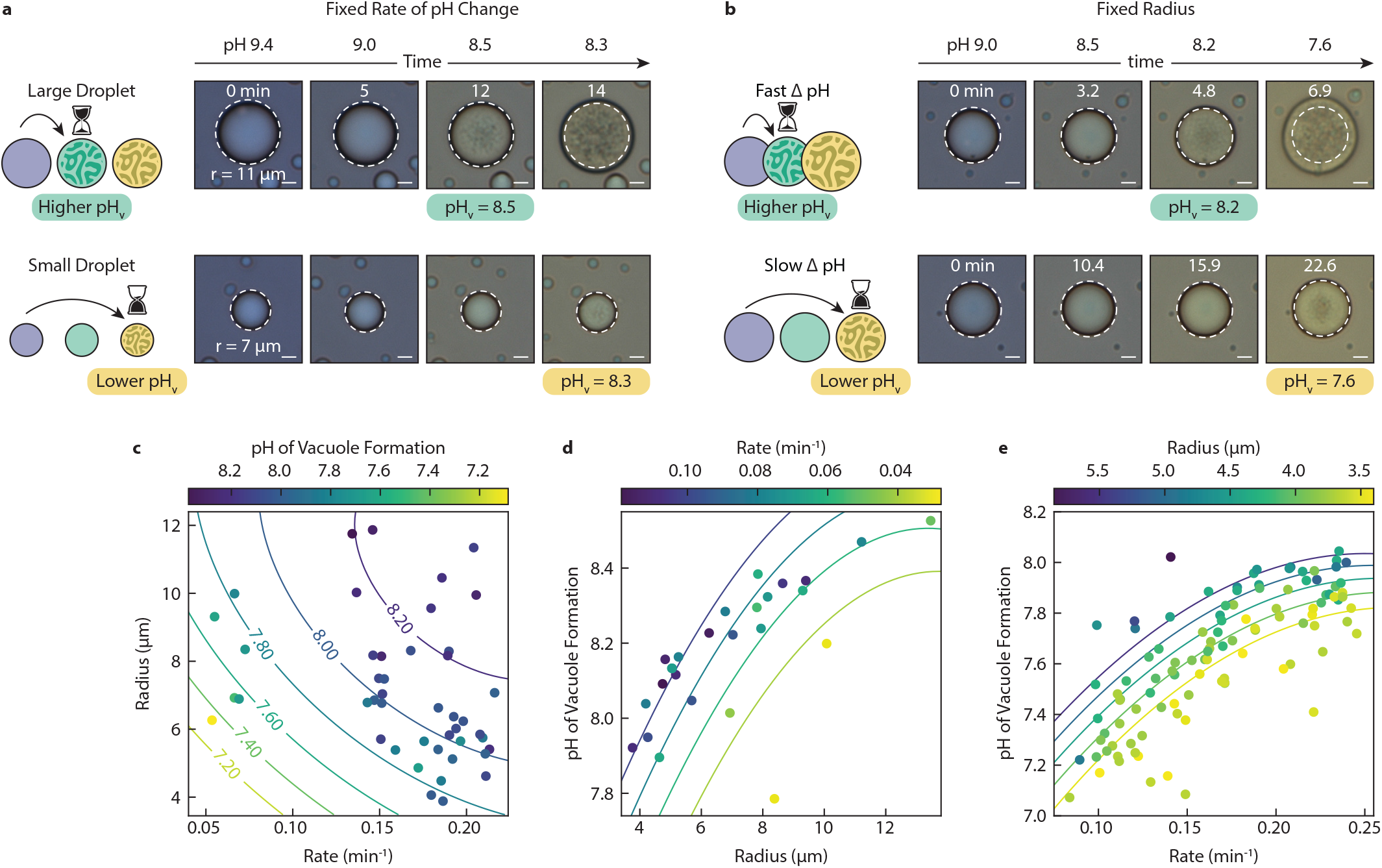
The pH of vacuole formation depends on the size of droplets and the rate of pH change. (a) Representative microscopy images of a large and small droplet over time as the pH decreases at the same rate. The larger droplet forms vacuoles earlier and at a higher pH. (b) Representative microscopy images of droplets of similar size over time as the pH decreases at varied rates. Vacuoles form depending on the rate of pH change, where a faster rate of change leads to earlier vacuole formation at a higher pH. (c) The pH of vacuole formation depends on both the droplet radius and the rate of pH change. (d) The pH of vacuole formation increases with the droplet radius within a given range of rates. (e) The pH of vacuole formation increases with the rate within a given range of drop radii. Data points represent droplets from a single capillary experiment in each of (c)-(e). Trend lines were created by linear regression to a parabolic surface.

The emergence of vacuoles is also a function of the rate of pH change (Fig. 3b). Imaging at different distances from the gel front allows drops exposed to different rates of pH change to be imaged in a single experiment (Fig. 3e). When controlling for drop size, faster rates of pH decrease result in vacuole formation at a higher pH. Comparing two drops of the same size (~ 9.4 *µ*m at pH 9.0), we see that the drop subjected to a faster rate of pH decrease (pH 9.0 to 7.6 over 7 min, rate = 0.174 *min*^−1^) exhibits the first signs of vacuole formation at pH 8.2 (Fig. 3b). In contrast, a droplet subjected to a slower pH decrease (the same drop in pH occurring over ~ 4 h, rate = 0.054 *min*^−1^) did not exhibit internal vacuole formation until the surrounding pH reached 7.6.

These results show that the onset of vacuole formation (characterized by pH_v_) is a function of droplet size and the rate of pH change. Imaging drops of varying sizes at increasing distances from the acid can capture both these dependencies in a single capillary experiment (Fig. 3c). In addition to vacuole formation, we observed that the shrinking of droplets prior to vacuole formation is also size and rate dependent. We hypothesize that the shrinking of drops may be due to the efflux of water and biomolecules as the drop composition changes in response to the external pH change. We compare the decrease in drop size among drops of varying sizes exposed to similar rates of pH decrease (Supplementary Fig. 10). We observe that larger drops shrink to a lesser extent than smaller drops when subjected to the same change in pH at the same rate. Similarly, the shrinking of coacervate drops of similar sizes is dependent on the rate of pH decrease (Supplementary Fig. 11). For a given pH change, coacervate drops exposed to slower rates of pH change shrink more than coacervate drops of similar sizes that experience faster rates of pH decrease. These observations imply that vacuole formation and the preceding reduction in coacervate drop size are non-equilibrium processes. At faster rates of pH decrease or in larger drops, the diffusion of polyelectrolytes across the drop boundary becomes the rate-limiting step, leading to vacuole formation and a smaller reduction in drop size. In contrast, at slower rates of pH decrease or within smaller drops, the drop has sufficient time to equilbrate with the surrounding pH changes, preventing vacuole formation and resulting in greater shrinkage of the drop.

### Vacuole formation requires rates of pH change faster than macromolecule relaxation

Our results show that the emergence of vacuoles in coacervate drops exposed to a pH decrease occurs when the exchange of components between the two phases becomes diffusion-limited. At a constant enzyme-to-polycation ratio within the capillary (3.16:1 enzyme-to-polymer), the equilibrium composition of each phase changes as a function of the pH. Initially homogeneous at pH 10, the droplets respond to a pH decrease caused by an acid front, equilibrating to the new conditions via polyion transfer. Our results show that larger droplets exposed to faster pH decreases are more prone to vacuole formation but less susceptible to volume reduction. This dependency suggests that the time scale for the diffusion of polyelectrolytes across the droplet boundary is slow compared to the rate of pH decrease at these conditions. To validate this hypothesis, we perform FRAP (fluorescence recovery after photobleaching) measurements to evaluate the mobility of the polycation within the condensate drop.

FRAP experiments show that the mobility of the polycation is relatively slow and has a similar time-scale as the formation of vacuoles (Fig. 4a). By bleaching a portion of a droplet containing FITC-labeled polycation, we can subsequently observe mobility of the polycation into the bleached region. The fluorescence recovery is fit to a biexponential with two time constants, with the slow time constant likely corresponding to dissociation of the polycation from the network. The slow recovery time (12 min) is similar to the time-scale for vacuole formation (3-50 min), and is likely artificially faster given the fluorescently labeled polymer used in these experiements was a lower molecular weight and therefore a lower net charge.

**Figure 4:**
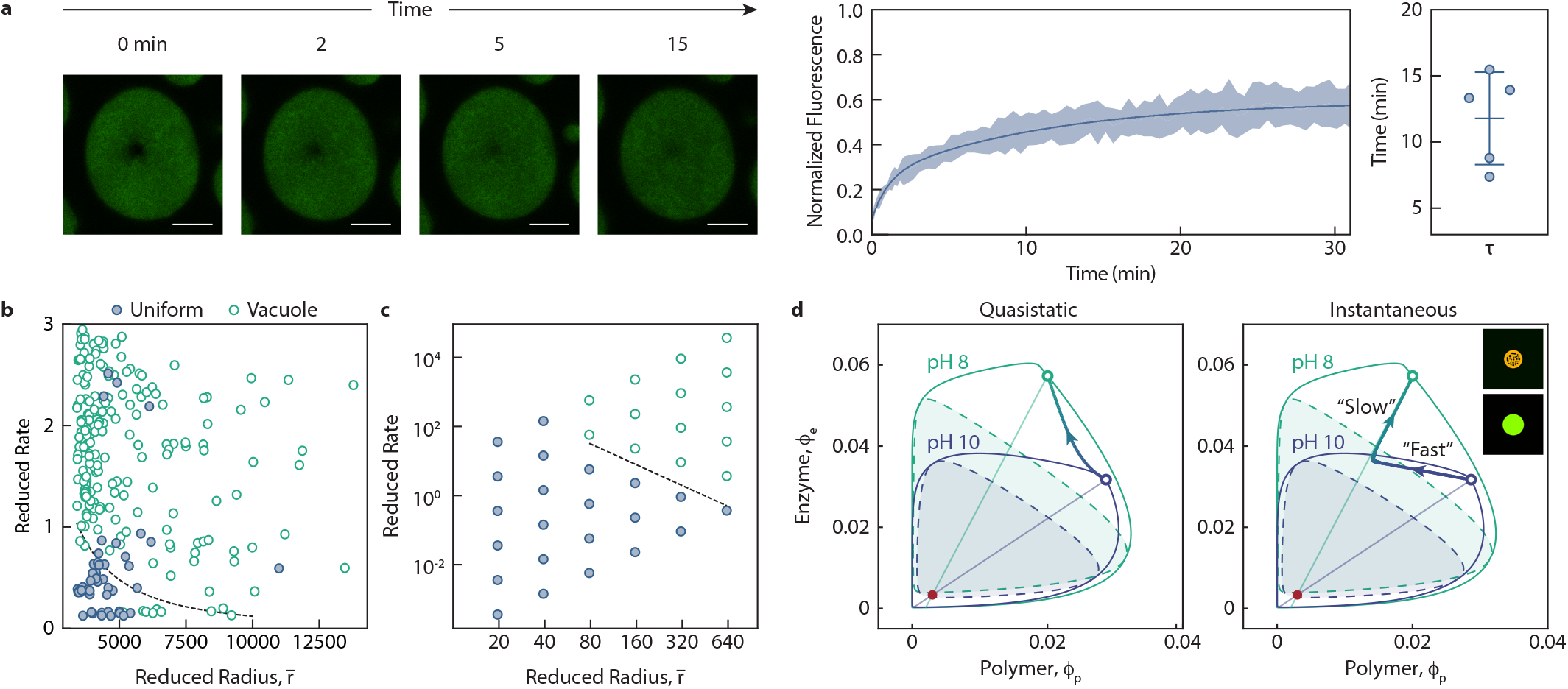
Slow mobility of polycation during a rapid pH change allows for vacuole formation. (a) FRAP experiments performed on coacervate drops show that the recovery of the bleached polymer occurs over 30 min. Confocal images of the bleached coacervate drop containing unlabeled enzyme and FITC labeled DEAE-dextran at 0.76 mass fraction of glucose oxidase in 10 mM Tris pH 9 buffer with 20 mM salt (1 mg/mL total macromolecular concentration). Scale bar 10 *µ*m. Biexponential fit (solid line) to the normalized recovery curves of five bleached drops. The shaded region represents ± 1 standard deviation from the mean intensity. The longer time scale for recovery of the bleached component was determined using the biexponential fit. Error bars indicate ± 1 standard deviation. (b) Vacuoles do not form in droplets if the rate of pH change is sufficiently slow and and the drops are sufficiently small. (c) Simulations demonstrate that sufficiently small droplets do not form vacuoles, regardless of the rate of pH change. The simulations also show a similar dependence of vacuole formation on the rate of pH change as in (b). (d) Ternary phase diagrams at pH 10 and pH 8 as predicted by the model with trajectories for quasistatic (left) and instantaneous (right) pH changes showing that fast rates of pH change result in trajectories that enter the spinodal region. Inset shows snapshot images from 2D simulations following an instantaneous pH change from pH 10 (bottom) to pH 8 (top).

The competition between the time scale of pH change and polycation transfer across the drop substantiates that if the rate of pH change is sufficiently slow or drops sufficiently small, then vacuoles should not form (Fig. 4b). This is further supported by analysis of drops from all capillary experiments, where drops that experience a pH drop down to 7 do not always form vacuoles despite a pH change from 10 to 7. These results indicate that drops can remain at equilibrium (or near equilibrium) with relatively slow changes, preventing vacuole formation.

### Vacuole formation is a non-equilibrium process plausibly resulting from spinodal decomposition

Our experimental evidence suggests that vacuole formation in this system is a non-equilibrium process. We propose that rapid system perturbations shift the phase boundary, driving the system into the spinodal region, where spinodal decomposition leads to vacuole formation in the droplets. Under equilibrium conditions, the energetic cost of forming additional interfaces makes such vacuole formation thermodynamically unfavorable. This supports the claim that vacuole formation is tied non-equilibrium mechanisms.

We developed a minimal theoretical model that captures how a phase separating system responds to changes in system conditions, particularly pH. Rapid changes in pH shift the phase diagram leading to spinodal decomposition inducing vacuole formation. The minimal model employs Flory–Huggins theory to describe a ternary mixture of polymer, enzyme, and water, incorporating pH-dependent interaction parameters *χ*_*ij*_. These *χ*_*ij*_ values serve as binary interaction parameters for each pair of components (polymer–enzyme, polymer–water, and enzyme–water). Experimental measurements inform the estimation of these parameters, revealing a binodal region that expands with decreasing pH (Fig. 4d).

The ternary phase diagrams predicted by the model at pH 10 and pH 8 is shown in Fig. 4d. The solid blue line denotes the binodal boundary at pH 10, and the dashed blue line denotes the corresponding spinodal. Likewise, the solid green line represents the binodal at pH 8, and the dashed green line represents the associated spinodal. The tie-lines (shown as blue and green straight lines) at pH 10 and pH 8, respectively, intersect at a common point (marked in red) that indicates the overall volume fraction of polymer and enzyme. A mixture prepared at this overall composition separates into macromolecule-rich and macromolecule-lean phases, situated on opposite sides of the tie-line at equilibrium. Thus, with a “quasistatic” pH change from 10 to 8, the system remains on the binodal curve (as shown by the quasistatic trajectory in Fig. 4d), leading to homogeneous droplets without vacuoles. These observations corroborate the notion that rapid changes in the system’s parameters are necessary for vacuole formation to occur.

To capture how sufficiently fast pH changes induce vacuole formation, we use the Cahn–Hilliard model to describe the spatiotemporal evolution of the polymer–enzyme–water system. Any changes in pH shift the phase diagram and drive the system to reorganize its composition and relax to the new equilibrium. The model shows two diffusive modes of relaxation (*λ*_1_, *λ*_2_) in the system, where *λ*_1_ corresponds to the *slow* mode and *λ*_2_ corresponds to the *fast* mode. The *fast* mode pulls the local composition towards the tie-line at rate *λ*_2_(*π/R*)^2^, while the *slow* mode moves the system along the tie-line towards the binodal at rate *λ*_1_(*π/R*)^2^. Here, *R* is the radius of the droplet undergoing pH perturbation. Under an “instantaneous” pH change, a separation in timescales between these two modes causes the dense-phase composition to relax along a two-part trajectory with a characteristic “elbow”, as shown in Fig. 4d.

This two-step relaxation trajectory can bring the system into the spinodal region, where the instability grows at a finite rate *γ*\*. If *γ*\* is greater than the *slow* rate *λ*_1_(*π/R*)^2^ than vacuoles form before the system moves out of the spinodal region. Since the *slow* mode scales with 1*/R*^2^ and *γ*\* is not a function of *R*, larger droplets are more prone to vacuole formation because they remain in the unstable region longer than smaller droplets. This suggests that droplets with larger radii and undergoing fast pH change should form vacuoles as seen in the experiments.

We solved the model numerically for a range of droplet sizes and rates of pH change to map out the dependence of vacoule formation on these two parameters. The droplets are subject to a logistic pH profile that has a characteristic rate *r*. We define a dimensionless rate parameter, *reduced rate*, as *r*/[*λ*_1_(*π/R*)^2^] and a dimensionless radius, *reduced radius*, as *R/l*, where *l* is the interfacial length of the droplet. The black dashed line shown in Fig. 4d indicates a boundary in parameter space separating droplets that form vacuoles from those that remain homogeneous. The slope reflects the 1*/R*^2^ dependence of the slow diffusive mode.

Simulation results show that for droplets larger than a critical radius, increased rates of pH change result in vacuole formation, whereas slower changes in pH result in uniform droplets (Fig. 4c). To connect with experiments, we set *l* ≈ 1 nm based on the enzyme size, which should have the same order of magnitude as the interfacial length scale. Rates are scaled by the inverse of the slow FRAP timescale, representing the slowest diffusive timescale measured experimentally. These choices align the simulation results with experimental data (Fig 4b), thereby demonstrating that the model accurately reproduces both the qualitative and quantitative behavior of vacuole formation across a range of droplet sizes and pH change rates.

## Discussion

Our results demonstrate that vacuole formation in coacervate droplets is a non-equilibrium process caused by the diffusion-limited exchange of polyions across the droplet boundary during rapid pH changes. A qualitative model supports these experimental findings, showing that the mechanism of vacuole appearance corresponds to spinodal decomposition. Vacuole formation occurs when the coacervate system enters the unstable region and instability growth is faster than the time required for the perturbed system to relax to its new equilibrium state via macromolecular diffusion.

We show how a droplet subject to a pH change relaxes to the new equilibrium via a twostep process with *fast* and *slow* modes of relaxation. The *fast* mode pulls the local composition towards the tie-line where, the enzyme-to-polymer ratio remains approximately constant indicating complexation between enzyme and polymer. The *slow* mode moves the system along the tie-line towards the binodal, capturing the collective diffusion of the polymer–enzyme complex and ensuring the dense phase attains the equilibrium density. The thermodynamic force driving complexation is stronger than the thermodynamic force driving diffusion of the complex across the droplet interface. Both rates scale with 1*/R*^2^, so larger droplets relax more slowly. Another consequence of the model is that sufficiently small drops 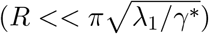 never form vacuoles, regardless of the rate of pH change because the slow mode of relaxation is still faster than *γ*\*. However, we could not validate this result experimentally.

This model provides a generalized framework for understanding vacuole formation in out of equilibrium associative phase-separated systems. Additionally, our capillary experimental setup, incorporating a novel colorimetric pH analysis method and detailed droplet characterization, offers a robust approach for studying vacuole dynamics in similar coacervate or condensate systems.

By advancing the mechanistic understanding and control over the internal organization of *in vitro* coacervate systems, our work provides insights into the behavior of *in vivo* hierarchical condensates and facilitates innovative applications in synthetic biology. Notably, the incorporation of an active enzyme offers the additional benefit of investigating vacuole formation within condensates driven by active processes, mimicking mechanisms observed *in vivo*.

## Methods

### Confocal experiments

Confocal images of the coacervate drops were taken using a Ti2 Nikon microscope with an AXR confocal unit in the Confocal and Specialized Microscopy Shared Resource (CSMSR) site at the Herbert Irving Comprehensive Cancer Center (HICCC). For imaging, coacervate drops were prepared using either glucose oxidase labeled with Alexa Fluor 594 NHS ester and unlabeled 500 kDa DEAE-dextran or unlabeled glucose oxidase and an 80:20 mixture of unlabeled 500 kDa DEAE-dextran and FITC-labeled 70 kDa DEAE-dextran (Millipore Sigma, product no. 54702) at a 0.76 mass fraction of glucose oxidase with a total macromolecule concentration of 1 mg/mL. Coacervate droplets were imaged in 10 mM tris buffer adjusted to pH 10 and supplemented with 20 mM NaCl. Droplets containing vacuoles were prepared via the addition of 0.1 M HCl to lower the solution pH prior to imaging.

### Passivation of imaging surfaces

Capillary surfaces were passivated with PEG to prevent interactions with the coacervate drops [17]. Capillaries were immersed in a 2 M NaOH solution for 30 min. They were then placed in 2 % (by volume) Hellmanex (Sigma Aldrich, product no. Z805939) solution in water and sonicated in a water bath for 10 min. The capillaries were rinsed with MilliQ water before submerging in ethanol and sonication in a water bath for 10 min. A fresh 5 mg/mL solution of mPEG-Silane (MW 5000, Laysan Bio) in ethanol with 1 % acetic acid was prepared by sonication in a water bath. Each capillary was filled with the mPEG-Silane solution and placed on a hot plate at 70 °C for 1 h. During this time, the capillaries were re-filled with the mPEG-Silane solution approximately every 15 min. PEGylation of CultureWell chambers (Grace Bio-Labs, product no. 112359) was carried out in a similar way with 200 *µ*L of mPEG-silane solution being added to each well at an interval of 30 min.

### Capillary experiments

A glucose oxidase solution (1 mg/mL) in water with 20 mM NaCl was prepared by dissolving the enzyme in MilliQ water and adjusting to pH *>* 8 with the addition of NaOH. The required amount of 5 M NaCl solution in water was added to the enzyme solution to bring the final salt concentration to 20 mM, followed by filtration with 0.2 *µ*m SCFA or PES syringe filters. A solutoin of 500 kDa DEAE-dextran (1 mg/mL) in water containing 20 mM NaCl was prepared in a similar way. A 20 mg/mL Xylenol Blue (Millipore Sigma 205486) solution in DMSO was prepared by adding a weighed amount of Xylenol Blue to DMSO and filtering the solution with a Nylon filter.To each of the enzyme and polymer solutions, the required volume of 20 mg/mL Xylenol Blue solution was added such that the final concentration of Xylenol Blue in each solution was 0.2 mg/mL. The pH of each solution was adjusted to 10 with 1 M NaOH and 1 M HCl using an InLab Pure Pro-ISM pH electrode. A 1 mL coacervate solution at 0.76 mass fraction of glucose oxidase was prepared by adding 0.76 mL of the glucose oxidase solution and 0.24 mL of the DEAE-dextran solution to an 1.7 mL tube and mixed by pipetting.

To prepare the gel loaded with acid, a 1% agarose solution in water (200 mg of agarose to 19.8 mL of MilliQ water) was heated in the microwave for 30 s. To the hot solution, 200 *µ*L of 20 mg/mL Xylenol Blue in DMSO and 20 *µ*L of 1 M HCl were added and mixed. A fixed amount of the hot liquid was pipetted onto a hydrophobic surface (plastic petri dish) and added to one end of the capillary tube by capillary action.

After waiting for a few minutes for the gel to solidify, the coacervate solution prepared as described above was added to the capillary tube using a syringe. The capillary was then secured to a microscopy slide on both ends using double-sided tape or a combination of hot glue and 5-minute epoxy. Time-lapse images at different distances from the gel interface were taken using the CMOS camera on the brightfield channel in an EVOS FL Auto 2 inverted fluorescent microscope.

## Model: Dynamics of vacuole formation

We model the phase diagram using Flory–Huggins theory for ternary mixture of polymer, enzyme, and water, incorporating pH-dependent interaction parameters *χ*_*ij*_.

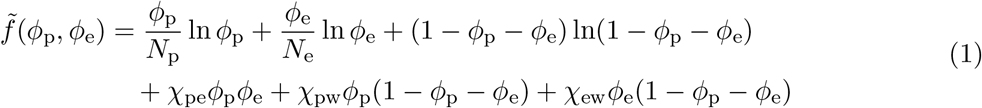

where *ϕ*_*i*_ is the volume fraction of species *i* = p, e, w such that Σ_*i*_ *ϕ*_*i*_ = 1, *k*_B_*T* is the thermal energy, *ν* is the volume per lattice site, and *χ*_*ij*_ are binary interaction parameters. The Flory-Huggins interaction parameters *χ*_*ij*_ are related to the pairwise interaction energies *u*_*ij*_ between components *i* and *j* as *χ*_*ij*_ = *z*(2*u*_*ij*_ − *u*_*ii*_ − *u*_*jj*_)*/*(2*k*_B_*T*), where *z* is the coordination number of the lattice.

Experiments suggest that the charge ratio between polymer and enzyme increases monotonically with decreasing pH. To capture electrostatic effects on the Flory–Huggins interaction parameters, *χ*_*ij*_ are represented as second-order functions of charge densities *ρ*_*i*_ and *ρ*_*j*_.

To incorporate the kinetics of phase separation we use the Cahn-Hilliard approach.

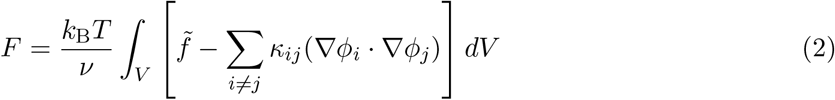

where 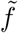 is the Flory-Huggins free energy density of equation (1), and the indices *i, j* take the values p, e, w corresponding to polymer, enzyme, and water, respectively. The parameters *κ*_*ij*_ control the energy and width of the interface between dense and dilute phases.

The local conservation of species *i* implies that its concentration, *c*_*i*_ = *ϕ*_*i*_*/ν*, evolves as

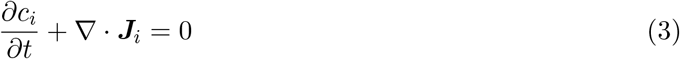

where ***J***_*i*_ are diffusive fluxes given by ***J***_*i*_ = − Σ_*j*_ *M*_*ij*_∇*µ*_*j*_ and *µ*_*i*_ = *δF/δc*_*i*_. The mobility coefficients *M*_*ij*_ depend on the local composition.

Within the bulk phase, the dynamic equations can be linearized about the homogeneous compositions 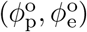. The linearized dynamics of the conserved field (*ϕ*_*p*_, *ϕ*_*e*_) is governed by

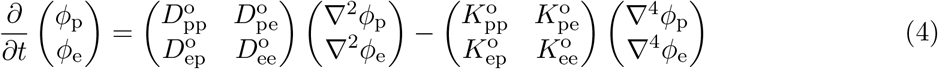

where the components of the diffusivity matrix ***D*** are given by the matrix product

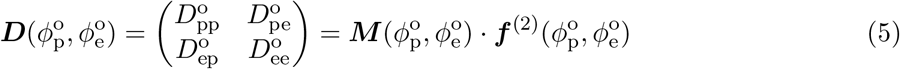

where 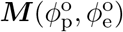 is the mobility matrix, and 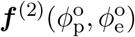 is the matrix of second derivatives of the free energy. Similarly, the coefficients of the ***K*** matrix are given by the matrix product

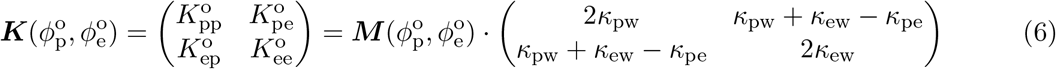

Notably, the diffusion matrix ***D*** has two approximately orthogonal eigenvectors (***e***_1_, ***e***_2_) with one vector parallel to the tie lines. The first eigenvector ***e***_1_, which is along the tie lines, corresponds to “slow” transport with diffusivity *λ*_1_. The second eigenvector ***e***_2_, which is approximately orthogonal to the tie lines, corresponds to “fast” transport with diffusivity *λ*_2_.

The model is solved numerically in one-dimensions and two-dimensions. The simulations start with a well-equilibrated uniform droplet at pH 10 in a periodic domain that is then subject to a logistic pH profile between pH 10 and pH 8. The imposed pH change is characterized by one single rate parameter *r*.

